# The helix bundle domain of primase RepB’ is required for dinucleotide formation and extension

**DOI:** 10.1101/2020.02.20.955914

**Authors:** Sofia Banchenko, Chris Weise, Erich Lanka, Wolfram Saenger, Sebastian Geibel

## Abstract

During DNA replication, primases synthesize oligonucleotide primers on single-stranded template DNA, which are then extended by DNA polymerases to synthesize a complementary DNA strand. Primase RepB’ of plasmid RSF1010 initiates DNA replication on two 40 nucleotide long inverted repeats, termed ssiA and ssiB, within the oriV of RSF1010. RepB’ consists of a catalytic domain and a helix bundle domain which are connected by long α-helix 6 and an unstructured linker. Previous work has demonstrated that RepB’ requires both domains for initiation of dsDNA synthesis in DNA replication assays. However, the precise functions of these two domains in primer synthesis have been unknown. Here we report that both domains of RepB’ are required to synthesizes a 10–12 nucleotide long DNA primer whereas the isolated domains are inactive. Mutational analysis of the catalytic domain indicates that the solvent-exposed W50 plays a critical role in resolving a hairpin structures formed by ssiA and ssiB. Three structurally conserved aspartates (D77, D78 and D134) of RepB’ catalyse the nucleotidyl transfer reaction. Mutations on the helix bundle domain are identified that either reduce the primer length to a dinucleotide (R285A) or abolish primer synthesis (D238A) indicating that the helix bundle domain is required to form and extend the initial dinucleotide synthesized by the catalytic domain.

## INTRODUCTION

Transfer of genetic information from viral, prokaryotic, eukaryotic and plasmid genomes to the daughter generation is ensured by the duplication (replication) of genomic DNA. During DNA replication primases synthesize *de novo* short oligonucleotide primers complementary to the single-stranded parental template DNA (1). DNA polymerases require the 3’-end of the primer for nucleotide addition, in order to synthesize a complementary DNA strand. Primers are elongated continuously on the leading strand and discontinuously on the oppositely oriented lagging strand resulting in the synthesis of Okazaki fragments, which require periodically primase activity.

The DNA replication of the broad host range RSF1010 has been extensively studied in *Escherichia coli*. Unlike its host genomes (i.e. *E. coli*) RSF1010 is replicated exclusively in leading strand mode (2, 3) (Supplementary figure 1a). RSF1010 encodes helicase RepA, primase RepB’ and initiator protein RepC, which are required for its own replication alongside the host replication machinery (4, 5). Binding of RepC to the origin of vegetative replication (oriV) initiates the unwinding of the double stranded plasmid DNA (dsDNA) by RepA in opposite directions (6, 7) and exposes two 40 nucleotide single strand initiator sequences A and B (ssiA, ssiB), which are encoded only once on complementary DNA strands (Supplementary figure 1b, c). RepB’ requires ssiA and ssiB to initiate DNA replication (2, 8, 9).

**Figure 1.**
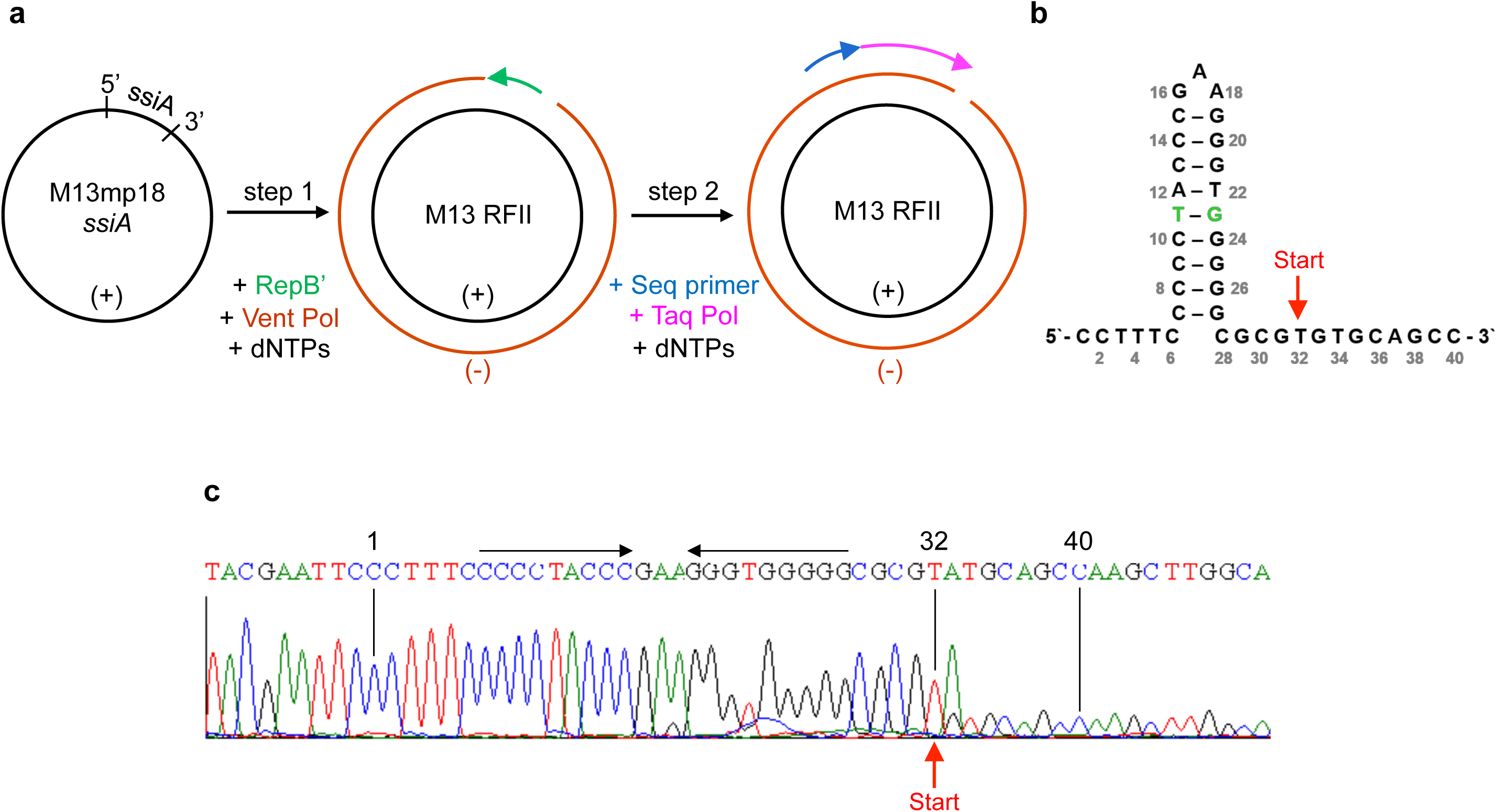
Primase RepB’ starts primer synthesis on thymine 32 of ssiA. **a)** Principle of the DNA replication assay. Primase activity is indicated by the conversion of single- to double-stranded DNA (step1). RepB’ synthesizes a complementary DNA primer (green arrow) on ssiA of the (+) strand of M13mp18ssiA ssDNA (black). The Vent DNA polymerase elongates the primer by addition of dNTPs complementary to the single-stranded M13mp18ssiA template (brown). The double-stranded DNA contains a single-strand nick and is labelled replicative form II (RFII), in order to distinguish it from the closed, circular M13dsDNA, which is labelled replicative form I (RFI). A *run off* DNA sequencing reaction was performed in order to find the first nucleotide of the RepB’ primer (step 2). B) Schematic of the ssiA hairpin. The starting point for the primer synthesis on thymine 32 is indicated with a red arrow. C) The sequencing signal drops at thymine 32. A mismatching adenine signal in position 33 of ssiA can be attributed to the Taq DNA polymerase, which adds dAMP to the 3’ end of DNA.

The unusually complex ssiA and ssiB are unique to IncQ and IncQ-like plasmids and exhibit hairpin loops at nucleotides 7-27 of ssiA/ssiB (10, 11)(Supplementary figure 1b and 1c). While RepB’ binds specifically the first six nucleotides of ssiA that lie upstream of the hairpin (10), *in vivo* studies suggested that primer synthesis may be initiated downstream of the hairpin within a triplet ‘GTG’ (nt 31-33) (2). *In vitro* DNA replication experiments showed that RepB’ requires dNTPs for primase activity.

Previous studies provided insights into the RepB’ primase mechanism. The X-ray structure of RepB’ exhibits a catalytic domain with a mixed α/β fold, that is connected via an unstructured linker to a helix bundle domain. The crystal structure of the catalytic domain of RepB’ bound to the first 27 nucleotides of ssiA revealed that a specific binding site for the first six nucleotides is in proximity to the active site suggesting that the missing nucleotides 28–40 of ssiA may bind along the catalytic centre and place the ‘GTG’ triplet (nt 31-33) close to the three catalytic aspartates 77, 78 and 134 (10). Comparisons of RepB’ with known structures of primases and polymerases revealed that these three aspartates are conserved, and may coordinate two metal ions for the catalysis of the nucleotidyltransferase reaction (12–14). DNA replication assays confirmed that RepB’ requires these three aspartates for the initiation of dsDNA synthesis (10). However, the catalytic domain and the helix bundle domain are inactive in isolation and DNA replication is only initiated in presence of both domains (10). The helix bundle domain of RepB’ binds ssiA with significantly lower affinity (∼ 27 µM) compared to the catalytic domain (∼ 2 µM), indicating a more transient interaction with ssiA compared to the catalytic domain (10). Therefore, it was proposed that the catalytic domain remains bound to the 5’-end of ssiA/ssiB during primer synthesis while the helix bundle domain stabilizes the growing primer on the DNA template (10).

Both, catalytic and helix bundle domain of RepB’ have structural orthologues in the archaeo/eukaryotic primase class despite the lack of sequence conservation (15). X-ray crystallography and biochemical studies of heterotrimeric primase PriSXL from *Sulfolobus solfataricus* revealed that helix bundle subunit PriX provides the nucleotide binding site for primer initiation while catalytic subunit PriS harbours the nucleotide binding site for nucleotide addition (16). As observed for RepB’, both subunits are far apart from each other in PriSXL indicating that the complex needs to transition into a closed conformation that juxtaposes the nucleoside triphosphates bound to initiation and catalytic sites for dinucleotide synthesis and subsequent elongation to synthesize a primer.

A nuclear magnetic resonance study on the primase part of primase-polymerase ORF904 from archaeal plasmid pRN1 showed the helix bundle domain of ORF904 bound to template DNA and two ATP molecules, suggesting that this structure represents an early step of the catalytic cycle before the template DNA and the two nucleotides are transferred to the catalytic domain for dinucleotide formation (17).

Several studies have shown that primases act as caliper to limit the primer length (18–20). Primases are classified into procaryotic DnaG type and archeal/eucaryotic Pri type primases. Although structurally distinct, both classes have in common that they require a catalytic and an accessory domain for primer synthesis. The latter one is a helix bundle domain in Pri type primase and a zinc binding domain in DnaG type primases, where it has been shown to initiate and regulate primer synthesis and length. The structural requirements that restrict the primer length remain elusive.

While most primases have no sequence specificity or recognize only triplets on template DNA, ssiA and ssiB are unusually long and complex and may have additional structural requirements for primer synthesis. Here, we set out to dissect the structure of RepB’ by analysing its primer synthesis in combination with *in vitro* DNA replication assays.

## RESULTS

### RepB’ begins primer synthesis on thymine 32 of ssiA

As starting point and length of the RepB’ primer were unknown, we sought to investigate the primer synthesis of RepB’ first. An *in vivo* study suggested that RepB’ begins the primer synthesis within nucleotides 31-33 of ssiA and ssiB (2). To verify the exact starting point of the primer synthesis, we established a DNA replication assay using purified RepB’ and Vent DNA polymerase, 2′-deoxynucleoside-5′-triphosphates (dNTPs) and single-stranded DNA of the phage M13mp18 (M13, Max-Planck strain 18) encoding ssiA in the (+) strand (M13mp18ssiA) (Figure 1a.b; Supplementary figure 2a,b). This reaction produced a dsDNA (M13 RFII, replicative form II) containing a nick in the newly synthesized (-) DNA strand where the primer synthesis was started (Figure 1a, step1). Next, we carried out a *run off* sequencing reaction towards the ssDNA nick using a sequencing primer complementary to the nicked (-) strand of the M13 RFII DNA (Figure 1a, step2). The fluorescence signal of the sequencing reaction fell off significantly after thymine in position 32 of ssiA showing that the first nucleotide in the RepB’ primer is a dATP (Figure 1c). An additional mis-matching adenine signal in position 33, that overlaps with the lower fluorescence signal of guanine 33 can be attributed to the Taq polymerase used in the sequencing reaction, which adds adenosine monophosphate to free 3’-OH termini independent of the DNA template.

**Figure 2.**
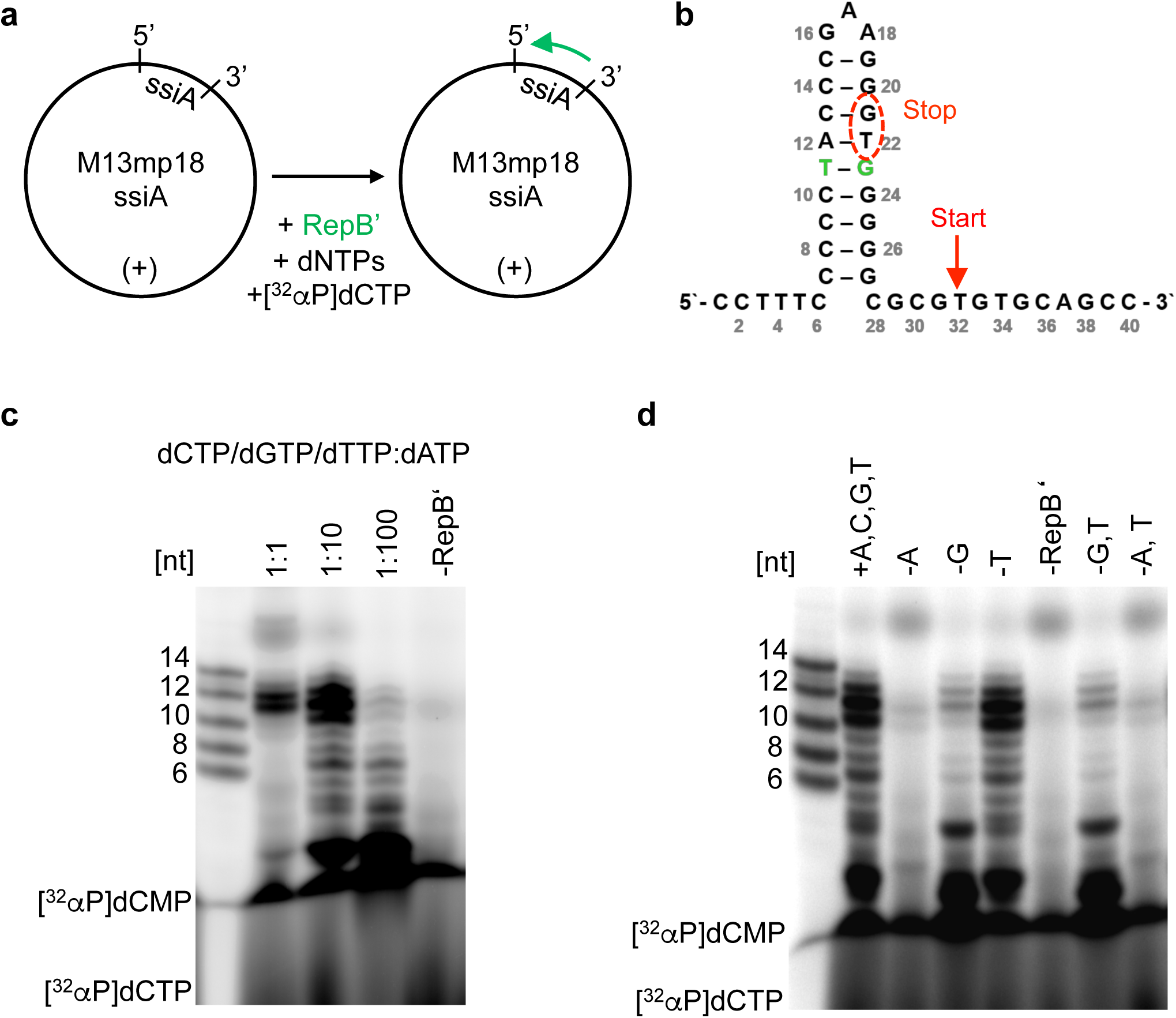
Primase RepB’ synthesizes 11 to 12 nucleotide long primers. **a)** Schematic of the primer synthesis reaction. **b)** Schematic of the ssiA sequence. Starting (red arrow) and end point (red circle) of the RepB’ primer are indicated. The wobble base pair is shown in green. **c)** DNA primer synthesis of RepB’ on circular single-stranded M13mp18ssiA. Oligonucleotide length standard: 6, 8, 10, 12 und 14 nucleotides (lane 1). Reactions contained M13mp18ssiA, RepB’ and 10 µM dCTP (1 µM [^32^αP] dCTP + 9µM dCTP), dGTP and dTTP. The dATP concentration was varied from 10 µM (lane 2), 1 mM (lane 3) to 10 mM (lane 4). The negative control contained all four dNTPs (10µM), M13mp18*ssiA* DNA, but no RepB’ (lane 5). The primer length was either determined by counting premature primer abortions or by comparison to the oligonucleotide marker. **d)** DNA primer synthesis of RepB’ on circular single stranded M13mp18*ssiA* DNA using combinations of dNTPs. Reactions contained RepB’, M13mp18ssiA and [^32^αP] dCTP, but different compositions of dNTPs. For maximal signal, dATP was used at 1mM and dCTP, dGTP and dTTP at 100µM concentration. Lane 1 oligonucleotide marker 6-8-10-12-14 nucleotides; Lane 2 dATP, dCTP, dGTP, dTTP; Lane 3 dCTP, dGTP, dTTP; Lane 4 dATP, dCTP, dTTP; Lane 5 dATP, dCTP, dGTP; Lane6 no RepB’ in the reaction; Lane 7 dATP, dCTP; Lane 8 dCTP, dGTP.

### RepB’ synthesizes DNA primers of up to 12 nucleotides

To investigate the primer synthesis, RepB’ was incubated with a mix of dNTPs supplemented with [^32^αP] dCTP and M13mp18ssiA (Figure 2a,b). Reactions were analysed on a denaturing polyacrylamide gel. At equimolar concentrations of all four dNTPs, two primers with a length of 11 and 12 nucleotides were observed (Figure 2c, lane 2). As dATP is the first nucleotide to be incorporated, we increased the dATP concentration 10-100 times in relation to dCTP, dGTP und dTTP (100µM each).

Higher dATP concentrations stimulated the primer synthesis considerably and led to the synthesis of shorter primers (Figure 2c, lanes 3 and 4). At a concentration of 1 mM dATP, RepB’ produced predominantly primers of 2, 10, 11 and 12 nucleotides while at a concentration of 10 mM dATP all [^32^αP] dCTP was incorporated into di-, tri- and hexa oligonucleotides. The continuous synthesis of short oligonucleotides requires dissociation of these primers from the M13mp18ssiA, which then becomes available as ssDNA template again. This result further indicates that RepB’ contains a nucleotide binding site that preferentially binds dATP. Once dATP is bound, synthesis of a new primer is initiated. For the subsequent experiments, we used an excess of 1 mM ATP in the reaction mix as this produced the highest amounts of full-length primer.

To confirm dATP as starting nucleotide of the primer, we carried out primer synthesis experiments using different combinations of dNTP. Full primase activity was only retained in presence of dATP (Figure 2d, lane 2), where as no primers were detected in the reaction without dATP (Figure 2d, lane 3). In absence of dGTP, RepB’ synthesized mainly dinucleotides and to a lesser extent trinucleotides as well as DNA primers with a length of up to 12 nucleotides (Figure 2d, lane 4). The accumulation of trinucleotides in the reaction without dGTP was expected because position C30 of ssiA requires the incorporation of complementary dGMP (3rd position in the DNA primer). However, the synthesis of trinucleotides and longer DNA primers in absence of dGTP shows that RepB’ also incorporates non-complementary dNTPs into the DNA primer, albeit inefficiently. Therefore, it is likely that in positions C28 and C30 of ssiA, dAMP has been incorporated into the RepB’ DNA primer to form a C-A wobble base pair. Omission of dTTP had no effect on the primer length because adenine does not occur in the sequence of ssiA, which serves as a template for primer synthesis (Figure 2d, lane 5).

### Potential role of W50 in unwinding the ssiA hairpin

Primer length and starting point of the primer synthesis indicate that RepB’ must unwind the ssiA hairpin structure, in order to synthesize primers in full length because the first base pair (C7-G27) of the hairpin is formed six nucleotides upstream of the primer start site (Figure 2b). The crystal structure of the catalytic domain of RepB’ bound to the first 27 nucleotides of the ssiA hairpin (10) shows that the solvent-exposed tryptophan 50 interacts with the first base pair and may play a role in unwinding the hairpin structure (Supplementary figure 3).

**Figure 3.**
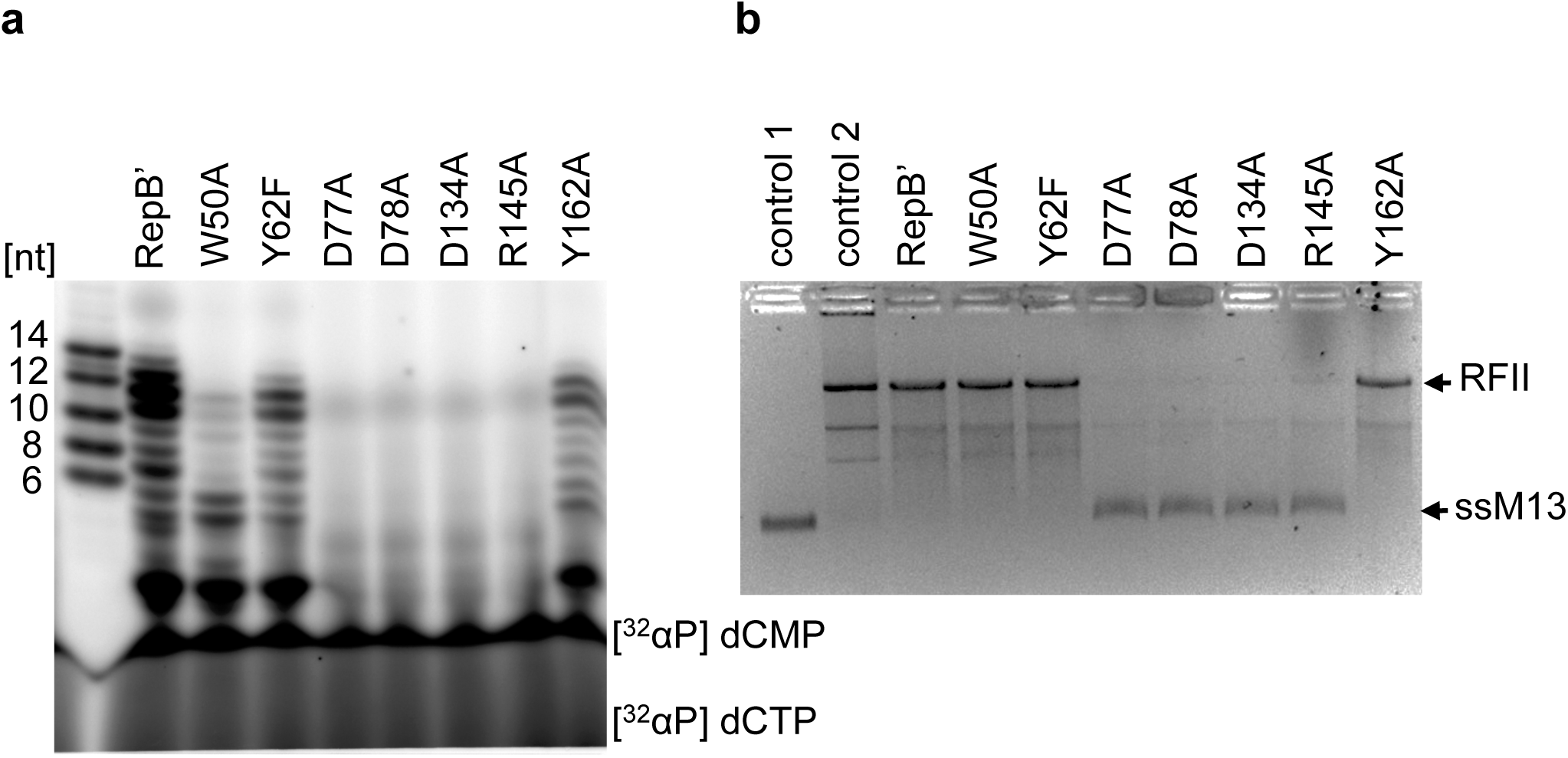
Primase activity of RepB’ variants W50A, Y62F, D77A, D78A, D134A, R145A, and Y162A. Possible role of W50 in hairpin unwinding. **a)** Primer synthesis assay. Reactions contained M13mp18ssiA, dNTPs, [^32^αP] dCTP and one of the following RepB’ mutants: W50A, Y62F, D77A, D78A, D134A, R145A, and Y162A. Oligonucleotide standard: 6, 8, 10, 12, 14 nucleotides. **b)** *In vitro* DNA replication assay. The assay tests the ability of the primase to initiate dsDNA synthesis on M13mp18ssiA in presence of the Vent DNA polymerase (see Figure 1A, step 1). Reactions contained Vent DNA polymerase, dNTPs, M13mp18ssiA and one of the following primase mutants: W50A, Y62F, D77A, D78A, D134A, R145A and Y162A. Controls: reaction without RepB’ (C1, lane 1), M13mp18*ssiA* dsDNA (RFI) (C2, lane 2).

To investigate whether W50 affects primer synthesis as well as the initiation of DNA replication, we used site-directed mutagenesis to generate mutant RepB’ W50A (Supplementary figure 2). We found that RepB’ W50A synthesized mostly short primers of 2, 5 and 6 nucleotides, whereas the synthesis of full-length primers was diminished significantly (Figure 3a). The predominant species of penta and hexa oligonucleotides indicates that primer synthesis is aborted before the hairpin or after breaking the first base pair of the hairpin pointing to a role of W50 in unwinding the ssiA hairpin. RepB’ W50A was still able to initiate DNA replication (Figure 3b).

### Three conserved aspartates and one arginine in the catalytic centre of RepB’ are required for nucleotidyltransferase activity

Comparison of the RepB’ with known primase and DNA polymerase structures revealed the spatially conserved amino acids D77, D78, D134 and R145, which form a nucleotide binding pocket on the catalytic domain (Supplementary figure 4). In order to investigate whether these amino acids are required for nucleotidyltransferase activity, primer synthesis of RepB’ variants D77A or D78A D134A and R145A was analysed on M13mp18ssiA (Figure 3B; Supplementary figure 2). The RepB’ mutants D77A or D78A or D134A did not synthesize detectable amounts of primers (Figure 3a) and – as reported previously (10) – could not initiate DNA replication (Figure 3b). We conclude that the amino acids D77, D78 and D134 catalyse the nucleotidyltransferase activity of RepB’. Although primer synthesis of RepB’ R145A was below the detection level, the mutant was still able to initiate DNA replication at a low level as previously shown (10), indicating basal nucleotidyltranferase activity below the detection level (Figure 3b). The strongly reduced primase activity of the RepB’ R145A suggests impaired binding or positioning of the incoming dNTP in the catalytic centre.

**Figure 4.**
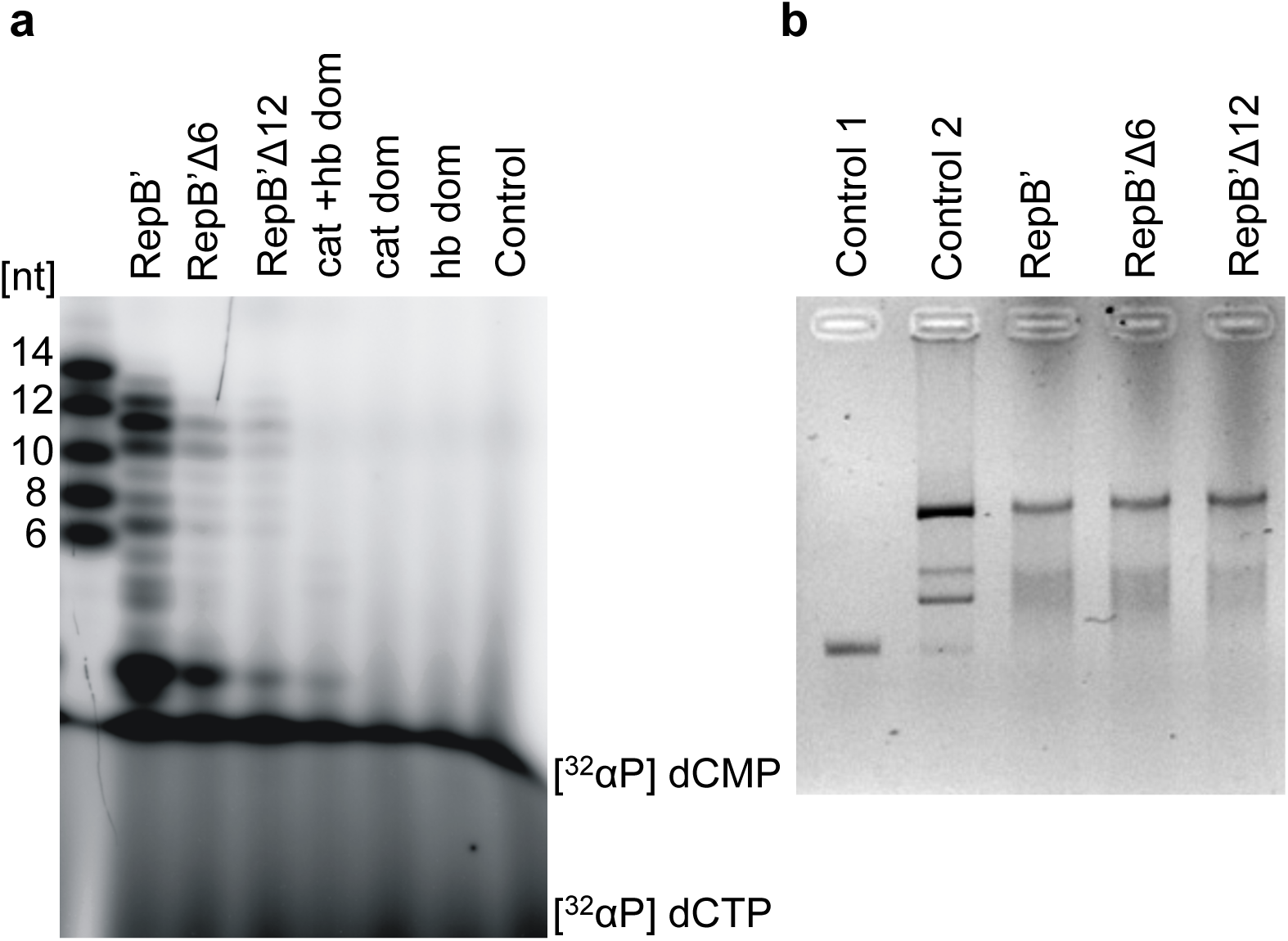
RepB’ requires catalytic and helix bundle domain for primer synthesis. **a)** Primer synthesis of RepB’Δ6, RepB’Δ12, (cat) catalytic and helix bundle domain of RepB’. The reactions contained M13mp18ssiA, dNTPs, [^32^αP] dCTP and one of the following RepB’ variants: RepB’Δ6, RepB’Δ12, catalytic domain (cat), helix bundle domain (hb), or cat + hb domains. Oligonucleotide marker (lane 1), control reaction without RepB’ (C, lane 8). **b)** *in vitro* DNA replication assay, representing the dsDNA synthesis by RepB’Δ6, and RepB’Δ12. Same controls are used like in the Figure 3 (C1, C2, RepB’).

Next, we investigated whether the two conserved amino acids Y62 and Y161, which are located on a flexible loop in proximity of the three catalytic aspartates, affect primase activity. Both RepB’ mutants retained full primase activity (Figure 3a, b; Supplementary figure 2).

### The helix bundle domain is essential for primer synthesis

Both, catalytic domain and helix bundle domain are required to initiate DNA replication (10). To investigate whether the helix bundle domain is involved in primer synthesis or required for handing over the synthesized primer to the DNA polymerase, we analysed primer synthesis of the catalytic (residues 1–212) and helix bundle domain (residues 213–323). When either of the two domains was tested in isolation, neither DNA replication (10) nor primer synthesis was detected (Figure 4a, lanes 6 and 7), whereas both domains combined, produced low amounts of di-, tri- and tetranucleotides (Figure 4a, lane 5). We conclude that the helix bundle domain is required for initiating primer synthesis. The lower primer synthesis of the separated domains compared to wild type RepB’ may be attributed to the missing linker, which imposes proximity of both domains.

### The linker connecting catalytic and helix bundle domains of RepB’ is required for full primase activity

To test how the linker length (residues 207–218) affects primase activity, we generated RepB’Δ6 and RepB’Δ12 with linker truncations of six (residues 207–212) and 12 amino acids (residues 207–218). Both RepB’Δ6 and RepB’Δ12, synthesized a lower amount of primers than wild type RepB’ (Figure 4a, lanes 3 and 4) and initiated DNA replication (Figure 4b, lanes 4 and 5). RepB’Δ6 synthesized higher amounts of primers than RepB’Δ12, indicating that the efficiency of primer synthesis depends also on the flexibility mediated by the linker between the two domains.

### The helix bundle domain is required for dinucleotide synthesis and primer elongation

To gain insights into the function of the helix bundle domain during primer synthesis, we sought to map functional amino acids on its surface. Structural comparisons with primases of the archeal/eukaryotic primase family revealed three conserved pockets with differential amino acid composition on the surface of the helix bundle domain of RepB’ (Supplementary figure 5a, b). In primase-polymerase ORF904, two of these pockets serve as binding sites for two ATPs whereas the other pocket binds template DNA. The comparison revealed only one invariant aspartate in the nucleotide binding pocket 1 of ORF904 (D308 ORF904, D238 RepB’), whereas no invariant amino acids were found in the nucleotide binding pocket 2 and the DNA template binding pocket. In ORF904 aspartate D308 is required for primer initiation. In PriX an arginine has replaced the aspartate in pocket 1 and the bound AMPNPP adopts a different orientation compared the ATP in ORF904. As we could not identify strictly conserved amino by structural comparisons, we carried out protein sequence comparisons of RepB’ with primases encoded by the RSF1010-related IncQ and IncQ-like plasmids (10). These comparisons revealed several conserved amino acids on the surface of the helix bundle domain. To validate their role in primer synthesis, an alanine scan of amino acids R234, D238, D281, R285 and E302, was performed. The RepB’ mutants R234A, D281A, and E302A synthesized reduced amounts of DNA primers compared to wild-type RepB’ (Figure 5a) and retained full primase activity in the replication assay (Figure 5b).

**Figure 5.**
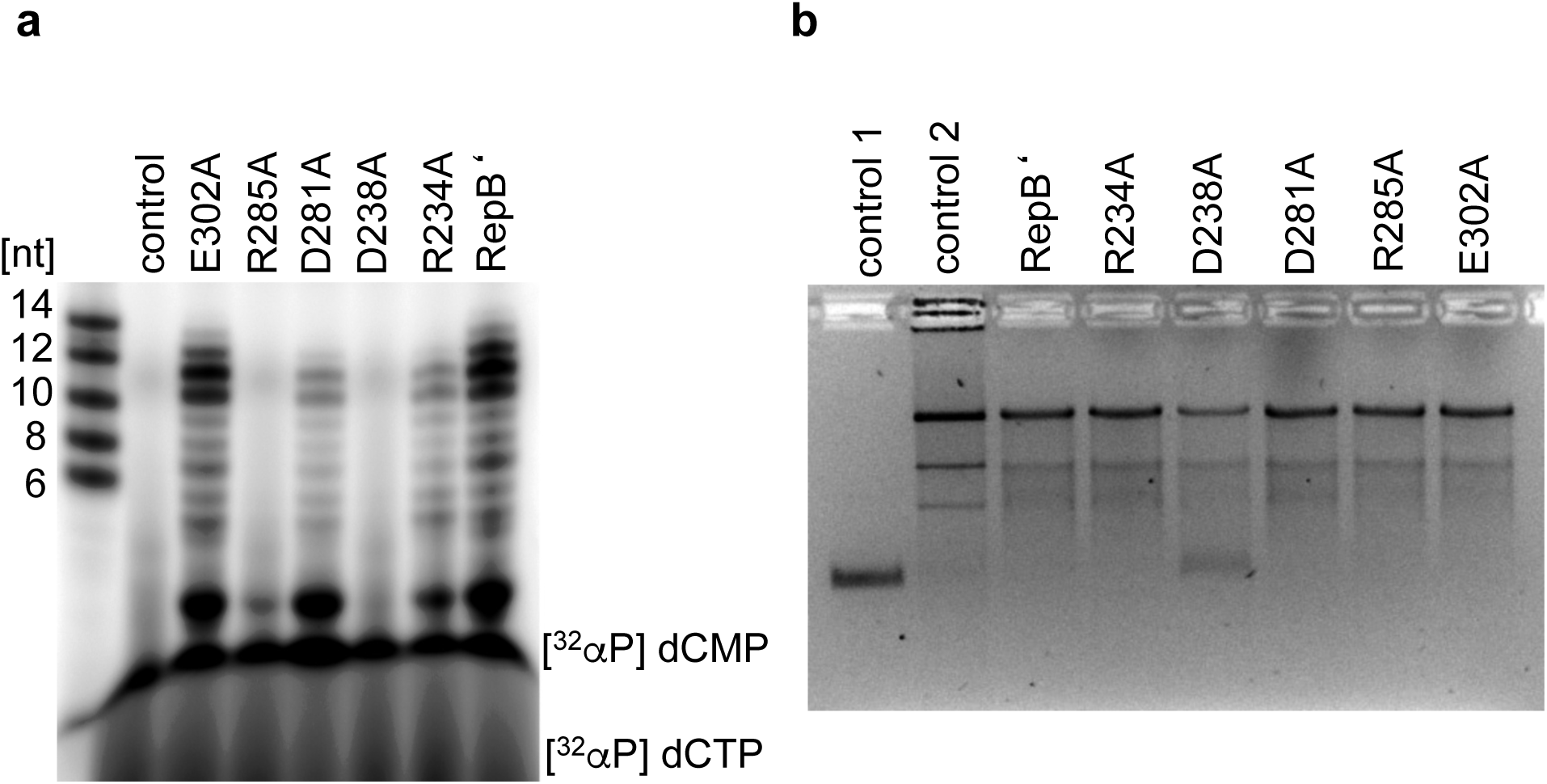
The helix bundle domain of RepB’ is required for dinucleotide formation and primer extension. Selected conserved amino acids on the helix bundle domain of RepB’ were mutated to alanine and their impact on the primer synthesis **(a)** and DNA replication **(b)** was tested. Primer synthesis of mutant D238A could not be detected while mutant R285A synthesizes a small amount of dinucleotides. However, both mutants, D238A and R385A, were able to initiate DNA replication.

In contrast, primer synthesis for RepB’ D238A was not detectable (Figure 5a, lane 6). This is in line with the conservation of D238 in nucleotide binding pocket 1 of ORF904. Thus, this result suggests that the helix bundle domain of RepB’ provides the second nucleotide binding site required for dinucleotide formation.

RepB’ R285A produced only low amounts of dinucleotides (Figure 5a, lane4). Longer primers were not detected suggesting that R285 is involved in the elongation step after dinucleotide formation. Interestingly, although not structurally conserved, R285 is placed into the DNA binding pocket of ORF904 in structural superimpositions. Furthermore, R285 binds a sulphate anion, suggesting that R285 could interact with the phosphate backbone of template DNA and/or the primer and stabilize the growing primer.

Despite the impaired primer synthesis both mutants, RepB’ D238A and RepB’ R285A, retained primase activity in the DNA replication assay (Figure 5b, lanes 5 and 7). The activity of RepB’ D238A was diminished compared to wild type RepB’. Taken together, these results show that the helix bundle domain is required for dinucleotide synthesis and primer elongation.

## DISCUSSION

Here we have investigated primase RepB’ of the eubacterial IncQ plasmid RSF1010, which is exclusively replicated in leading strand mode. Compared to known primases, RepB’ has two highly similar, unusually complex recognition sites. The catalytic domain of RepB’ binds the first six nucleotides of ssiA at the 5’ end before the hairpin structure (nt 7–27). Here, we have shown that RepB’ synthesizes an 11–12 nucleotide long DNA primer starting with dATP at thymine 32 on the 3’ end of ssiA. Our data indicates that W50 is involved in unwinding the hairpin during primer synthesis, which becomes necessary after the primer has exceeded a length of five nucleotides.

We show that the catalytic domain of RepB’ harbours a nucleotide binding site, which catalyses nucleotidyltransferase reaction. Furthermore, our mutational analysis of the helix bundle domain along with structural comparisons of archeal/eukaryotic primases indicate that the helix bundle domain of RepB’ has a dual function in binding ATP for primer initiation and in elongating the dinucleotide, possibly by stabilizing the initial dinucleotide primer/ssDNA template. Our results imply that RepB’ must transition into a closed active conformation, which brings the nucleotide binding sites of catalytic and helix bundle domain in proximity for the initial formation of a dinucleotide. We show that dinucleotide formation is considerably stimulated by dATP. Without dATP, primer synthesis is strongly reduced showing that the incorporation of the first nucleotide is base-specific and not dependent on the sugar or tri-phosphate moiety. We also observe premature abortion of primer synthesis in presence of excess dATP. Thus, the reoccupation of the dATP binding site triggers the formation of a new primer indicating that the binding sites for primer initiation and elongation stay close together during primer synthesis.

Our observations are in agreement with the general model for primer synthesis put forward by Charles Richardson (1). This model suggests that the primase catalyses the formation of a phosphodiester bond between two adjacent nucleotides. Primer synthesis requires the hydrolysis of the nucleotide at the elongation site. The triphosphate moiety of the nucleotide at the initiation site is preserved and this nucleotide becomes the 5’ end of the primer - in case of RepB’ dATP.

After dinucleotide formation RepB’ moves along the template DNA while nucleotides are being added to the 3’ hydroxyl group of the primer until an oligonucleotide primer of defined length is synthesized. Interestingly, we found that RepB’ variants with truncations in the linker still synthesize a full-length primer, although such truncations do not permit interaction between catalytic and helix bundle domain. This is because α-helix 6 acts as a spacer between the two domains in the RepB’ crystal structure. Thus, our data suggests that α-helix 6 must be repositioned upon activation of RepB’ to bring catalytic and helix bundle domain in proximity, whereas the unstructured linker provides the flexibility for optimal interaction of the helix bundle domain with the growing primer. A model for the primer synthesis of RepB’ is presented in supplementary figure 6.

## METHODS

### Cloning

In order to generate truncations of the linker between catalytic and helix bundle domain, we amplified the DNA fragments encoding the catalytic domain (fragment nt 1-618) and fragments nt 642–969 und nt 657–969 encoding the C-terminal part of the protein including linker and helix bundle domain, respectively (Supplementary Figure 2a). Fragment 1–618 was cloned into pET28b, using *Nde*I and *Eco*RI restriction enzyme sites. Fragments 639–969 und 657–969 were cleaved using *Eco*RI and *Hin*dIII first and then inserted into pET28b encoding fragment 1-618 via *Eco*RI and *Hin*dIII, respectively. The phenylalanine triplet, introduced by the *Eco*RI site, was removed using the QuikChange® Site-Directed Mutagenesis Kit (Agilent), respectively producing constructs RepB’Δ6 (207–212) and RepB’Δ12 (207–218). Site-directed mutagenesis was used to generate RepB’ mutants W50A, Y62F, Y162A, R234A, D238A, D281A, R285A und E302A using plasmids pET28b:repb’ as template and the primers listed in table 1.

**Table 1.**
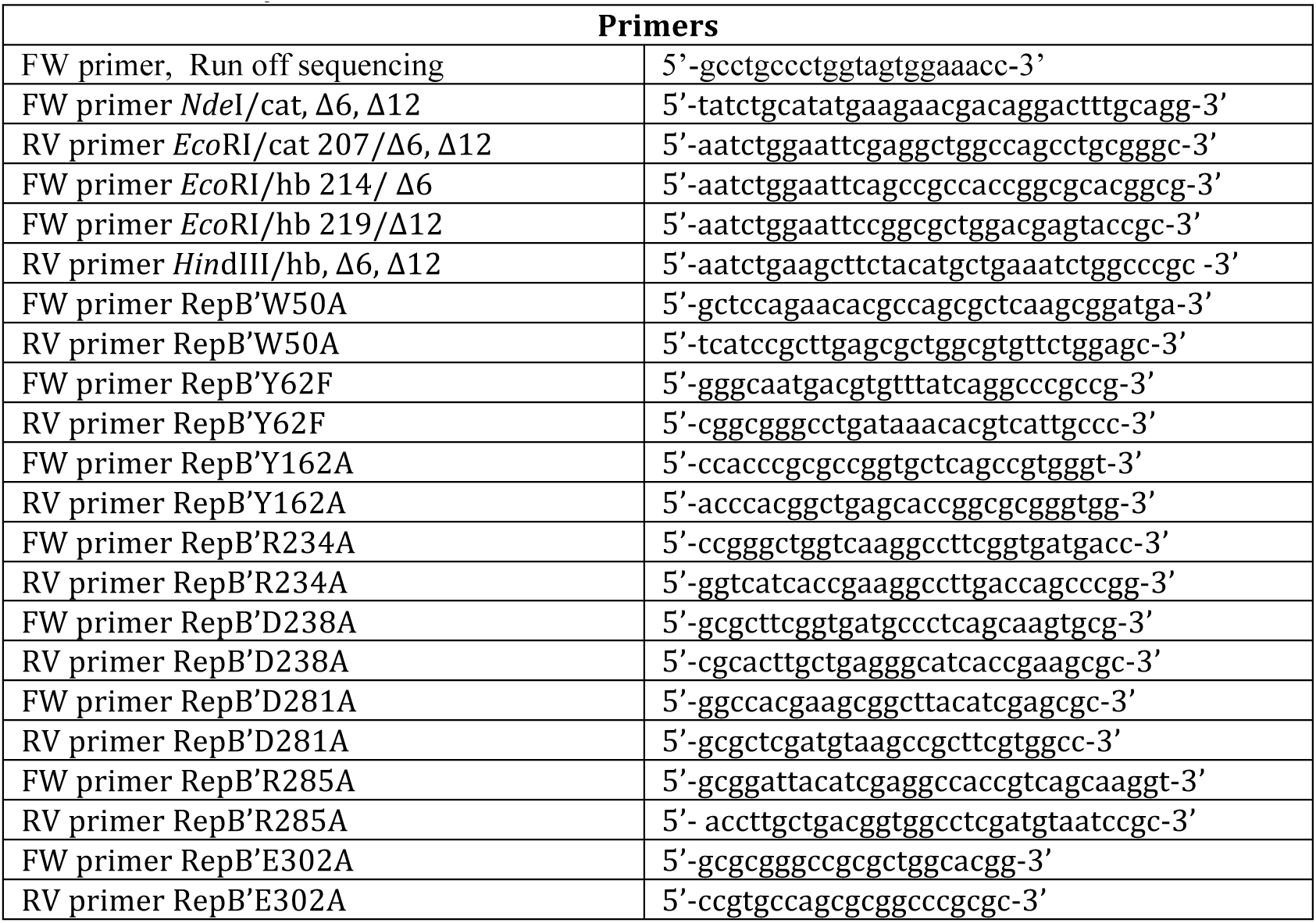
Primers used for cloning and mutagenesis. Point mutations and restriction enzyme sites are underlined.

### Protein expression and purification

RepB’ and its derivates W50A, Y62F, Y162A, R234A, D238A, D281A, R285A, E302A, RepB’Δ6 and RepB’Δ12 were expressed and purified using the published protocol for RepB’ (10) (table 2). *E. coli* ER2566 was transformed with the respective plasmids. Bacteria were grown in LB medium to an optical density of 0.6 at 600 nm (OD_600_). Overexpression was induced by addition of 0.4 mM IPTG (isopropyl β-D-1-thiogalactopyranoside). After 3 hours cells were harvested by centrifugation (10 min, 5,000 g, 4 °C) and resuspended in lysis buffer (50 mM Tris/HCl (pH 7), 150 mM MgCl_2_, 2.5 mM DTT). Cells were passed three times through a C5 EmulsiFlex homogenizer (Avestin). Cell debris was removed by centrifugation (30,000 g, 30 min, 4 °C), the supernatant diluted 1:2 with buffer A (20 mM Tris/HCl (pH 7), 50 mM MgSO_4_, 1 mM DTT) and then loaded on a HiTrap Heparin column (CV, column volume 5 ml; GE Healthcare), equilibrated with buffer A. The column was washed using a step gradient of 14 % buffer B (20 mM Tris/HCl (pH 7), 50 mM MgSO_4_, 2 M NaCl, 1 mM DTT), followed by application of a linear gradient (14-21 %, 5 CV). The eluted proteins were diluted 1:4 in buffer A and loaded on a 20 HS-column (CV 7 ml, Perseptive Biosystems), equilibrated with buffer A. Proteins were eluted over a linear gradient (0-20 %, 5 CV) of buffer B. The purest fractions were pooled and separated on a HiLoad^TM^-16/60-Superdex75 column (GE Healthcare), equilibrated with buffer C (20 mM Tris/HCl (pH 7), 150 mM MgSO_4_, 1 mM DTT). All proteins were analyzed by the SDS-PAGE to confirm their purity (Supplementary Figure 2b).

**Table 2.** Constructs used in this study.

| Construct | Reference |
| --- | --- |
| RepB' (1-323, full length) | (10) |
| RepB' catalytic domain (1-206) | (10) |
| RepB' helix bundle domain (206-323) | (10) |
| RepB' $\Delta 6$ ($\Delta$ 207-212) | This study |
| RepB' $\Delta 12$ ($\Delta$ 207-218) | This study |
| RepB'W50A | This study |
| RepB'Y62F | This study |
| RepB'D77A | (10) |
| RepB'D78A | (10) |
| RepB'D134A | (10) |
| RepB'R145A | (10) |
| RepB'Y162A | This study |
| RepB'R234A | This study |
| RepB'D238A | This study |
| RepB'D281A | This study |
| RepB'R285A | This study |
| RepB'E302A | This study |

The catalytic domain and the helix-bundle domain were expressed and purified as described in (10). Briefly, the two His-tagged proteins were expressed in *E. coli* ER 2566 (NEB) under the same conditions as described above for RepB’. Cell lysates were loaded on HisTrap column (CV 5 ml; GE Healthcare) equilibrated with buffer D (20 mM Hepes (pH 8), 200 mM MgCl_2_, 1 mM DTT) and eluted over a linear gradient of 0-50 % (100 ml) buffer E (20 mM Hepes (pH 8), 200 mM MgCl_2_, 1 M imidazol, 1 mM DTT). The His-tag was cleaved off using thrombin and removed by dialysis against buffer C. The catalytic domain was further purified using the RepB’ purification protocol for cation exchange and size-exclusion chromatography (20 HS & S75) as described above (Supplementary Figure 3).

### Production of circular single- and double-stranded DNA of phage M13mp18ssiA

The recombinant phage M13mp18ssiA (mp18, Max-Planck strain 18), which carries the ssiA sequence in the multi cloning site, was generated in a previous study (10). To produce ssDNA and dsDNA of M13mp18ssiA, electro competent *E. coli* strain JM103 carrying the F’-plasmid were transformed with M13mp18ssiA dsDNA and were grown at 37 °C in 400 µl LB medium for one hour. A volume of 100 µl indicator bacterium *E. coli* JM103 and 400 µl *E. coli* transformed with M13mp18ssiA dsDNA were mixed with 3 ml of 0.7 % soft agar and spread on X-Gal/IPTG agar plates containing 100 µg/ml ampicillin and incubated at 37 °C until blue pseudo plaques appeared on the bacterial loan. A phage suspension was prepared by inoculation of 3 ml LBM medium (LB medium, supplemented with 5 mM MgCl_2_) with a well separated pseudo plaque. In order to produce M13mp18*ssiA* dsDNA, a 3 ml pre-culture of indicator bacterium *E. coli* JM103 was grown in LBM medium at 37 °C for 3 hours (250 rpm) and then diluted 1:10 with LBM medium in a 250 ml Erlenmeyer flask (250 rpm). The 30 ml main culture was then infected with 3ml phage suspension. The infected culture was grown for 5 hours at 37 °C (500 rpm). Bacteria were harvested (5 min, 16,000 g, 4 °C) and the M13mp18ssiA RF-DNA extracted.

To produce M13mp18ssiA ssDNA, a pre-culture of *E. coli* JM103 (M13mp18ssiA^+^) was diluted 1:50 in TY medium. A volume of 6 ml diluted culture were mixed with 1.5 ml phage suspension and incubated for 5 hours at 37 °C in a 250 ml Erlenmeyer flask under vigorous shaking (450 rpm). Bacteria were pelleted by centrifugation (5 min, 16,000 g, 4 °C) and the phage containing supernatant transferred into a 2 ml Eppendorf tube and centrifuged again (5 min, 16,000 g, 4 °C) to remove residual bacteria. Finally, a volume of 1 ml of the phage suspension was transferred into a new 1.5 ml Eppendorf tube, mixed with 200 µl ice cold PEG-NaCl solution (20 % PEG 6000, 2,5 M NaCl) and precipitated on ice for an hour. The phages were pelleted (5 min, 16,000 g, 4 °C) and resuspended in 10 µl M13-Puffer (20 mM Tris/HCl (pH 7,6), 10 mM NaCl, 0.1 mM EDTA).

The phage suspension was heated to 60 °C and mixed thoroughly with 50 µl hot phenol (60 °C). The mixture was cooled down to RT, vortexed and centrifuged (3 min, 16,000 g). The aqueous phase was taken off, mixed with 100 µl chloroform, centrifuged (16,000 g, RT) and the supernatant transferred into a 1.5 ml Eppendorf tube. The DNA was mixed with 1/10 volume of Na-acetate (3 M, pH 5.2) and 2.5 volumes ethanol (100%), precipitated for 2 hours at −20 °C. After centrifugation (15 min, 16,000 g, 4 °C), the supernatant was discarded and the DNA pellet washed with 70 % ethanol, centrifuged (1 min, 16,000 g, 4 °C and the washing step repeated with 100% ice cold ethanol. The DNA pellet was dried at RT, resolved in 100 µl TS buffer (1 mM Tris/HCl, (pH 8.0), 1 mM NaCl), heated for 10 min at 65 °C) and cooled down to RT.

### DNA replication assay

The initiation of complementary DNA strand synthesis was tested *in vitro* using a published protocol (10). Briefly, 0.5 units Vent DNA polymerase, 50 nM primase RepB’, 30 ng M13mp18 ssDNA, 375 µM dNTPs were incubated in 1xNEB buffer for 10 min at 37 °C followed by an 8 min incubation time at 72 °C. The reaction was stopped by addition of 0.5 % SDS. Samples were separated by electrophoresis on 1 % TAE agarose gels and the DNA stained with 0.5 % ethidium bromide.

### Primer synthesis assay

The reaction contained 3 µM primase, 240 ng M13mp18ssiA ssDNA, 10 µM dCTP, 90 µM [^32^αP] dCTP (3,000 Ci/mmol), 100 µM dGTP, 100 µM dTTP, 1 mM dATP in 1xNEB-2 buffer. After incubation for 1 hour at 37 °C, formaldehyde loading buffer (95 % formaldehyde, 18 mM EDTA, 0.025 % SDS, 0.25 % xylene cyanol, 0.25 % bromophenol blue) was added and the heated for 2 min at 95 °C to stop the reaction. Samples were cooled down on ice before separation by vertical electrophoresis under denaturing conditions on a 20 % polyacrylamide gel (28 cm x 16 cm x 0.2 cm), containing 8 M urea and 1 x TBE buffer (pH 8).

A phosphor-screen (Type MS, Perkin Elmer^TM^) was exposed to the gel over night at −80 °C and read out using a phosphor scanner (Cyclone, Perkine Elmer). Oligonucleotides of 6, 8, 10, 12, 14 nucleotides in length (TIBMOL BIOL, Berlin) were radioactively phosphorylated with [^32^γP] dATP at their 5’ ends using phosphokinase (NEB) following the manufacturer protocol and used as a length standards.

### Run off DNA sequencing reaction

A DNA replication assay was carried out as described. The DNA sequencing was carried out by AGOWA *Genomics* (Berlin). Primers are listed Table 1.

## Acknowledgement

We thank Claudia Alings and Clemens Langner for technical support in the lab, Dr. Eberhardt Scherzinger for advice.

This work was supported by the German Research Foundation (SA196/43-1) and the Elite Network of Bavaria (N-BM-2013-246).

## Supplementary Figures

**Figure S1.**
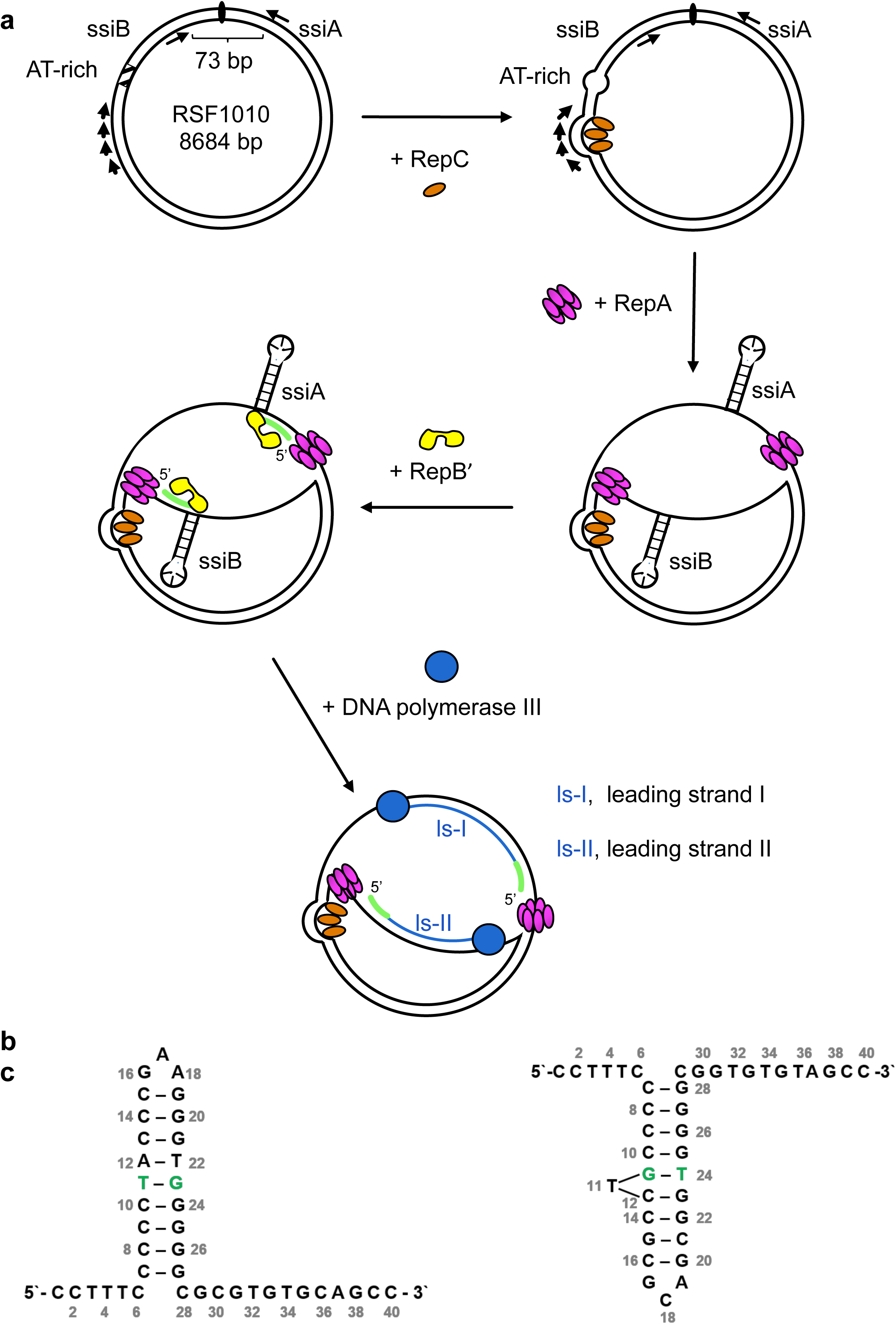
**a)** Replication of plasmid RSF1010. **b)** ssiA and **c)** ssiB form a hairpin structures (nt 7-27).

**Figure S2.**
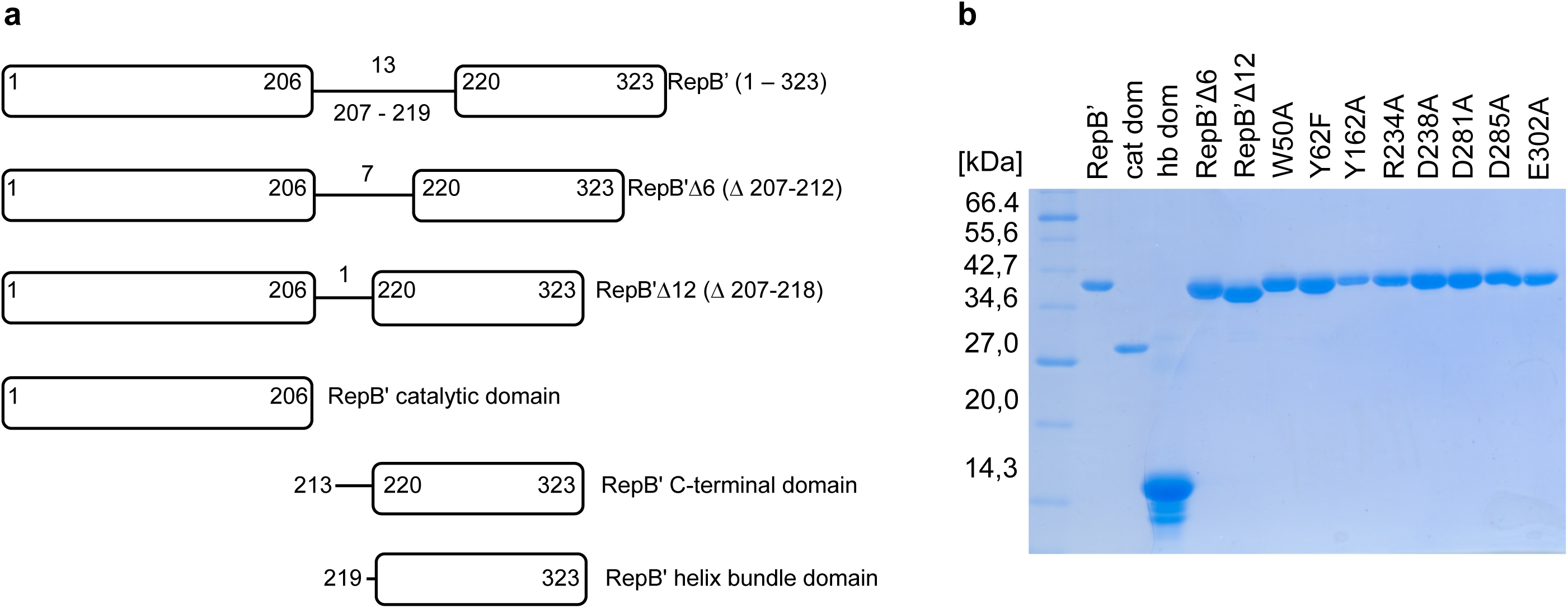
a) Schematic of RepB’ constructs used in this study **b)** SDS-PAGE of purified proteins used in this study. Lane 1: (1) P7702 protein molecular weight standard (NEB). Lane 2-14: RepB’, catatalytic domain (cat dom), helix bundle domain (hb dom), RepB’Δ6, RepB’Δ12, W50A, Y62F, Y162A, R234A, D238A, D281A, R285A, and D302A.

**Figure S3.**
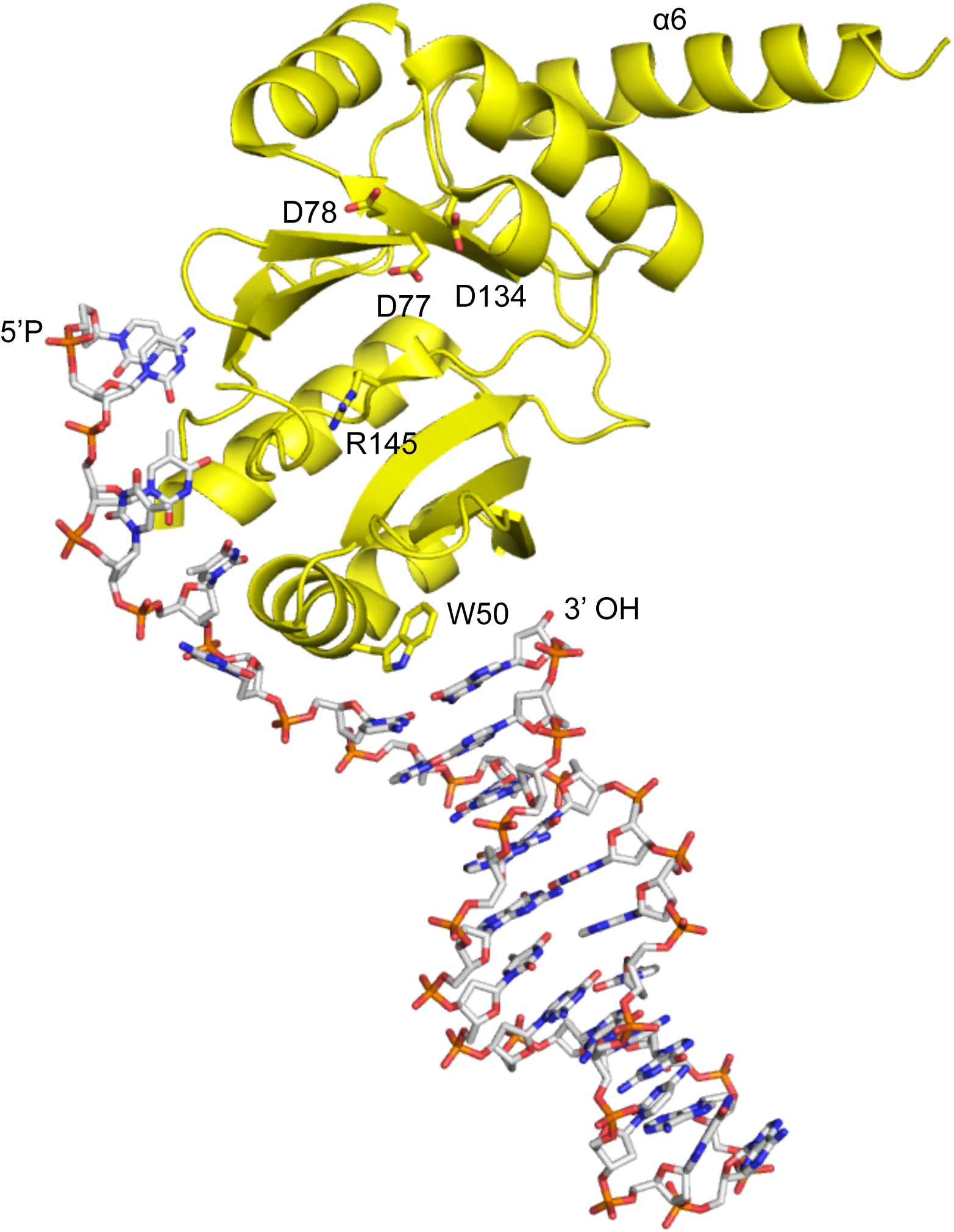
Crystal structure of the catalytic domain of RepB’ bound to the first 27 nucleotides of ssiA. Mutated amino acids shown as sticks.

**Figure S4.**
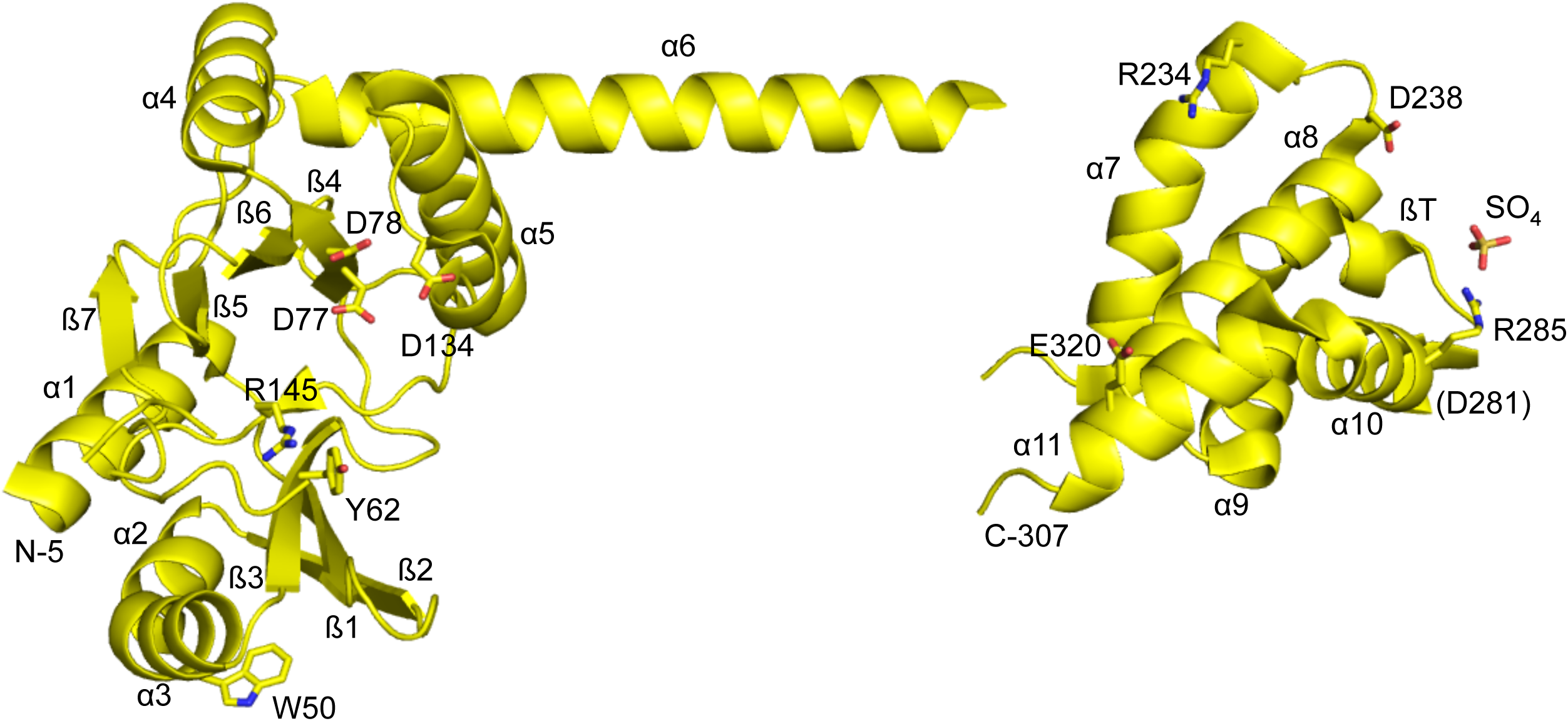
Crystal structure of RepB’. Mutated amino acids shown as sticks.

**Figure S5.**
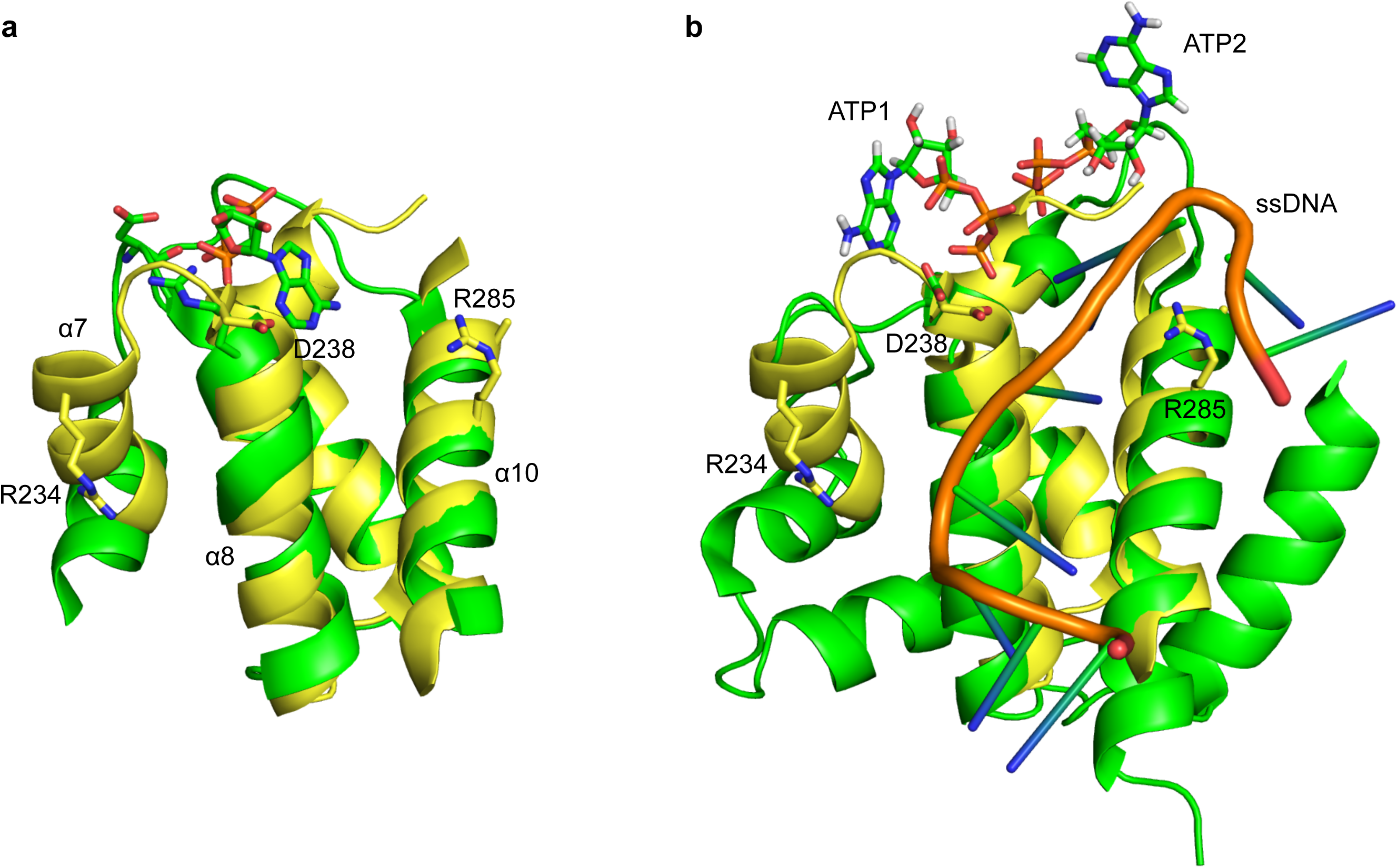
Superimposition of helix bundle domains of RepB’ (aa 225–296, yellow), ORF904 (green) and PriX (aa 58–126, green).

**Figure S6.**
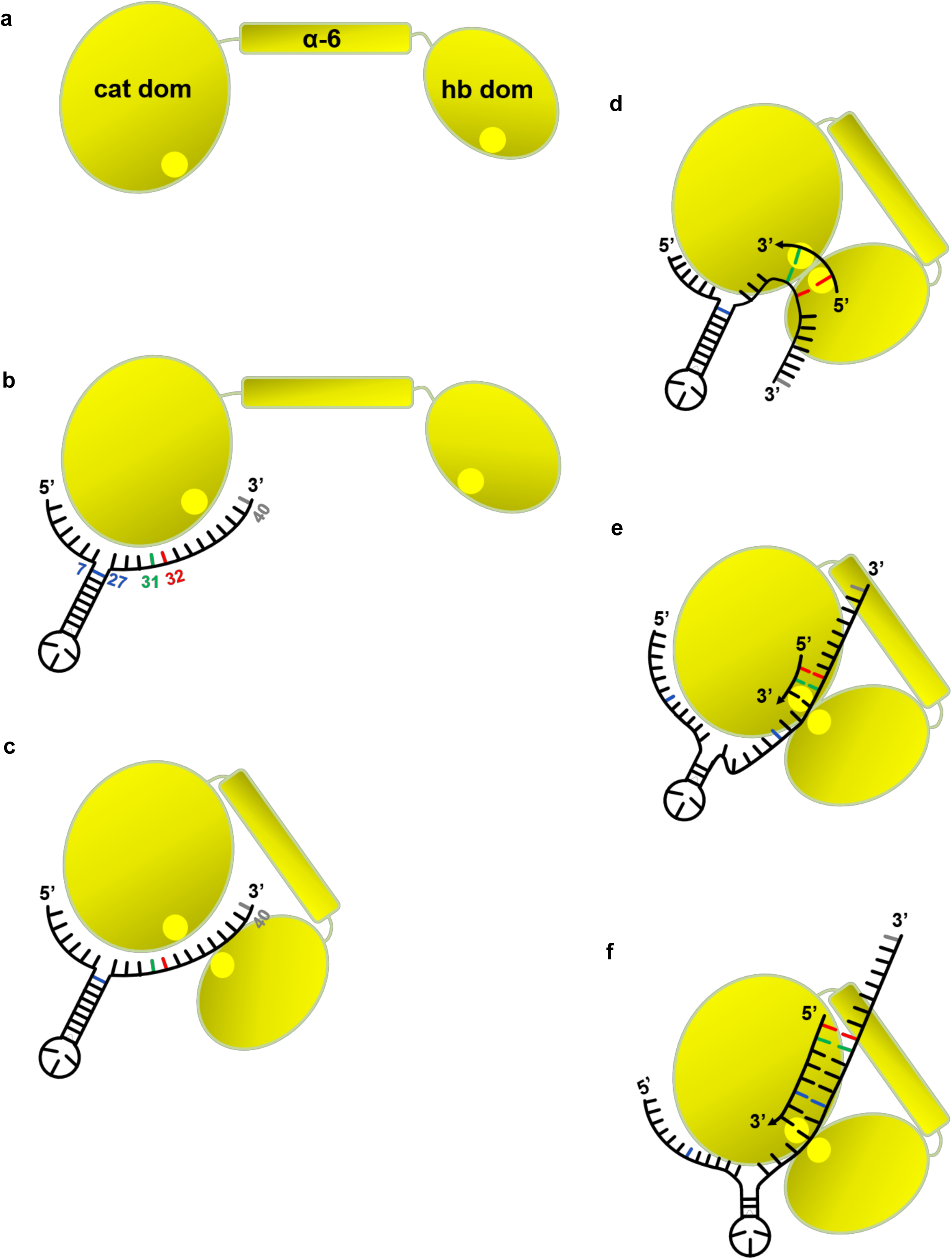
Schematic of the RepB’ primase mechanism.

